# Assessment of clonal kinetics reveals multiple trajectories of dendritic cell development

**DOI:** 10.1101/167635

**Authors:** Dawn Lin, Andrey Kan, Jerry Gao, Edmund Crampin, Philip D. Hodgkin, Shalin H. Naik

## Abstract

A thorough understanding of cellular development is incumbent on assessing the complexities of fate and kinetics of individual clones within a population. Here, we develop a system for robust periodical assessment of lineage outputs of thousands of transient clones and establishment of *bona fide* cellular trajectories. We appraise the development of dendritic cells (DCs) from barcode-labeled hematopoietic stem and progenitor cells (HSPCs) by serially measuring barcode signatures, and visualize this multidimensional data using novel developmental interpolated t-distributed stochastic neighborhood embedding (Di-SNE) time-lapse movies. We identify multiple cellular trajectories of DC development that are characterized by distinct fate bias and expansion kinetics, and determine that these are intrinsically programmed. We demonstrate that conventional DC and plasmacytoid DC trajectories are largely separated already at the HSPC stage. This framework allows systematic evaluation of clonal dynamics and can be applied to other steady-state or perturbed developmental systems.

## INTRODUCTION

Many conventional models of hematopoiesis consider a stepwise restriction of lineage commitment from hematopoietic stem cells (HSCs) to unipotent progenitor populations via multipotent and oligopotent stages (Guo et al., 2013; Månsson et al., 2007). Recent clonal fate (Ema et al., 2014; Lee et al., 2017; Naik et al., 2013; Notta et al., 2015; Sanjuan-Pla et al., 2013; Yamamoto et al., 2013a) and/or single cell RNA-sequencing studies (Nestorowa et al., 2016; Olsson et al., 2016; Velten et al., 2017) demonstrate that significant lineage imprinting is already in place within individual hematopoietic stem and progenitor cells (HSPCs) with branching occurring earlier than previously appreciated. However, most lineage tracing studies only measured clonal fate at a single time point. Therefore, questions remain as to whether the fate bias observed at one snapshot in time is consistent with earlier or later times, and therefore whether there may be asynchronous ‘waves’ of output over time.

Some studies have assessed clonal contribution longitudinally (e.g. by serially sampling progeny derived from HSCs in the blood) and have been instrumental in highlighting clonal properties including repopulation kinetics and lineage bias of HSPCs (Dykstra et al., 2007; Kim et al., 2014; Naik et al., 2013; Sun et al., 2014; Verovskaya et al., 2013; Wu et al., 2014; Yamamoto et al., 2013b). However, these approaches are only feasible i) for tissues that can be serially sampled from an animal, ii) for time points that are relatively far apart (e.g. weeks to months) and therefore restricted to long-term propagating stem/progenitor cells, and iii) for progeny that are sufficient in number to be sampled. Tracking clonal output longitudinally from transient progenitors, with low numbers of progeny and at higher frequency can benefit from long-term imaging, which allows accurate reconstruction of pedigrees. However, due to technical demands it generally only allows assessment of 10s-100s clones for a period of days to weeks (Skylaki et al., 2016). Recent ‘pedigree’ tools that measure evolving barcodes in progeny can infer developmental history (Frieda et al., 2016; Kalhor et al., 2017; McKenna et al., 2016), but are limited in their assessment of clonal kinetics.

Another method that aims to recapitulate the dynamic aspects of development and differentiation are ‘pseudo-time’ analyses, which infer developmental trajectories by assuming single cells within a population represent different ‘snapshots’ along archetypal paths, and align cells based on their proteomic or transcriptomic profiles (Wagner et al., 2016). These models can be of great benefit in understanding the order of gene/protein expression in developmental pseudo-time. A confounding factor, however, is the inability to assess individual clones as data is derived from a snapshot assessment, or with no lineage connection when assessed between time points. Therefore, such archetypal trajectories may mask heterogeneity at the clonal level, including features such as kinetics, lineage bias and division destiny (Marchingo et al., 2014).

Dendritic cells (DCs) represent a distinct branch of hematopoiesis and are responsible for pathogen sensing and activation of the adaptive immune response (Merad et al., 2013). There are three major subtypes including plasmacytoid DCs (pDCs), conventional DC type 1 (cDC1s) and cDC type 2 (cDC2s) (Guilliams et al., 2014). DC development is relatively well established at the population level, and can be recapitulated in cultures of bone marrow cells with *fms*-like tyrosine kinase 3 ligand (FL) (Naik et al., 2005). According to the current model, all DC subtypes can be generated from a restricted common DC progenitor (CDP) population downstream of HSPCs (Naik et al., 2013; 2007; Onai et al., 2007) via discrete subtype-committed precursor stages (Grajales-Reyes et al., 2015; Naik et al., 2006; Onai et al., 2013; Schlitzer et al., 2015). Importantly, clonal evidence has suggested earlier subtype-specific imprinting (Lee et al., 2017; Naik et al., 2007; Onai et al., 2007). However, as these clonal studies assessed fate at one snapshot in time, there is the possibility that fate was different at earlier or later time points. One study examined clonal DC development longitudinally using long-term imaging from CDPs (Dursun et al., 2016). However, the fate of many clones could not be accurately determined as they did not reach full differentiation at the end of the tracking period.

Cellular barcoding allows tracking of clonal fate by differential tagging of individual progenitors with unique and heritable DNA barcodes (Bystrykh et al., 2012; Naik et al., 2014). Quantification and barcode comparison between progeny cell types allows inference of lineage relationships i.e. barcodes shared between cell types infers common ancestors, whereas differing barcodes infers separate ancestors. Here, we combine cellular barcoding and DC development in FL cultures to facilitate longitudinal assessment of clonal kinetics in a robust, controlled and high throughput manner. Our results highlight that there are several distinct classes of cellular trajectories in DC development: each consist of clones with a similar pattern of DC subtypes produced over time but with varying properties including the timing, duration and magnitude of clonal waves. Importantly, using clone-splitting experiments, we demonstrate that many of these cellular trajectories are ‘programed’ within individual HSPCs. Furthermore, we demonstrate that pDC and cDC development has already largely diverged at the HSPC stage, and not downstream in the CDP as is currently assumed. Our results offer a powerful analytical and visualization framework that reveals the diversity of clonal kinetics and cellular trajectories.

## RESULTS

### Longitudinal tracking of clonal DC development reveals time-varying patterns

To track clonal DC development longitudinally, we barcode-labeled mouse Sca1^+^ cKit^hi^ cells that contained early HSPCs and cultured them with FL to allow DC generation (Figure 1A & S1A). The cultures were serially split in two at various times such that half of the cells were sorted for the DC subtypes using flow cytometry for subsequent barcode analysis, and half were kept in culture with a compensating amount of fresh media (Figure 1A). To accurately define pDCs, cDC1s and cDC2s, we used CD11c, MHCII, Siglec-H, CCR9, Sirpα and CD24 (Figure 1B). In addition, we sorted cells that were outside these DC gates collectively as “non-DCs” to allow estimation of the recovery of barcodes in the culture at any given time points and track clones that still contained DC progenitors. CCR9 inclusion was critical to define *bona fide* pDCs as Siglec-H^+^CCR9^−^ cells generated cDCs upon re-culture (Figure S2) (Schlitzer et al., 2011). Importantly, individual samples were separated into technical replicates after sorting and cell lysis to allow assessment of technical variation of barcode recovery (Figure 1A). Furthermore, control experiments (Figure S1B) were performed and demonstrated that serial sampling of barcoded progeny at the indicated time intervals was a robust approach to measure DC clonal kinetics (Figure S1C&D).

**Figure 1.**
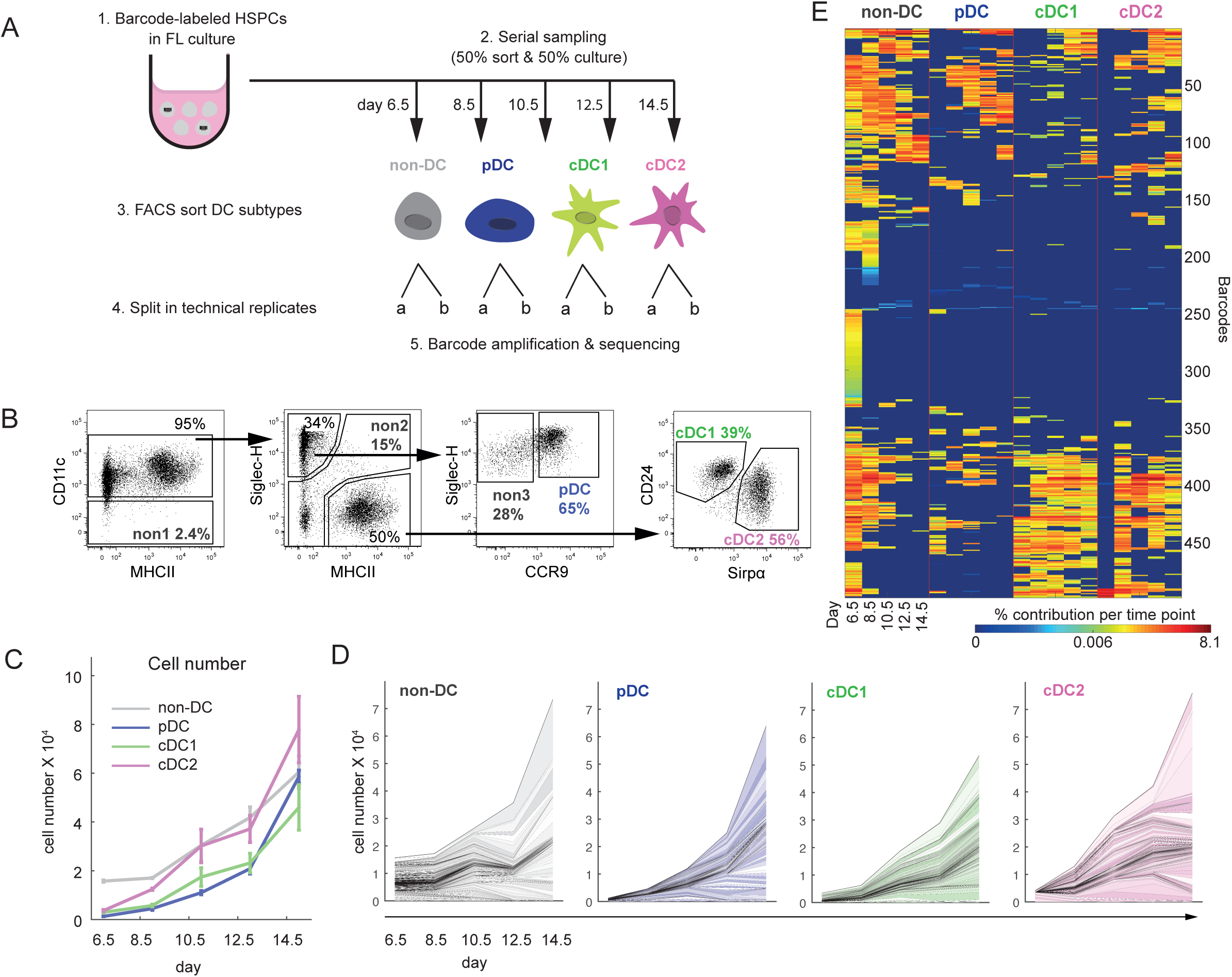
Longitudinal tracking reveals asynchronous waves of DC generation. (A) Experimental set up. HSPCs (cKit^+^Sca1^+^) from mouse BM were transduced with lentivirus containing DNA barcodes and cultured in FL-supplemented DC conditioned medium. At each time point, cells were equally split for either DC subtype isolation or further development in culture. non-DCs, pDCs, cDC1s and cDC2s were sorted as in (B) at each time point. Samples were then lysed and split into technical replicates, and barcodes were amplified and sequenced. (B) Gating strategy to isolate pDCs, cDC1s, cDC2s and non-DCs using CD11c, MHCII, Siglec-H, CCR9, Sirpα and CD24. Numbers represent % of cells from parent gate. (C) Number of DC subtype generation over time at the population level. (D) Stacked histogram showing clonal contribution (i.e. per barcode) to each DC subtype number over time. It is apparent that clones differ in size and also in timing of expansion. (E) Heatmap representation of clonal output to DC subtypes from individual time points. Data shown in C are average + SEM of 3 independent cultures from one experiment, representative of 3 independent experiments. D & E show all clones from one representative culture.

Our method assessed DC developmental dynamics and revealed novel time-varying patterns. First, we generated stacked histograms showing the number of cells produced by each clone at different time points (Figure 1C&D). We observed a temporal shift of DC contribution by a spectrum of large and small clones, and this pattern was apparent for all DC subtypes. This indicated that DC generation was not sustained by a set of ‘stable’ clones within the tracking period, and the contribution by different clones was not equal. We also generated a heatmap showing the barcode contribution to the number of DCs (biomass) from all cell types at all time points to capture the entirety of the data (Figure 1E). Again, the shift of clonal contribution to cell types over time was apparent, as was their bias.

Previous studies demonstrate that single HSPCs are biased towards a certain DC output (Lee et al., 2017; Naik et al., 2007; 2013). However, as fate was only assessed at a single time point, it is not clear how many DCs, and which subtypes of DCs, were generated at earlier or later times (i.e. in different parts of a putative clonal ‘wave’), which could lead to misclassification. For example, if a multipotent clone generated pDCs at an early time point and cDCs later, it would be classified as having a pDC-only or cDC-only fate depending on which time point was assessed. To test this, we first categorized clones into four classes (non-DC, pDC-only, cDC-only and pDC/cDC) and determined that only ∼30-40% of clones generated DCs when considered at any given single time point (Figure 2A & S1E). However, when we compared the ‘across time’ fate, taking into account a clone’s capacity to produce DCs at multiple time points, that proportion of DC-generating clones increased to nearly 90%. In addition, ∼20% of clones were re-classified from unipotent (pDC- or cDC-only) when measured at single time points to multipotent (pDC/cDC) when all time points were considered (Figure 2A). The asynchronous contribution to different DC subtypes over time was indeed apparent in the majority of clones using violin plots (Figure 2C). Therefore, fate should be considered in the context of time for a full appreciation of a clone’s potential. We further quantified the contribution to the number of DC subtypes by different classes of clones based on the definition ‘across time’ and observed lower contribution by multipotent (∼40%) than unipotent (∼60%) clones to both pDCs and cDCs (Figure 2B). These results highlight the importance of tracking development longitudinally to accurately and thoroughly interpret cellular output. Furthermore, our results indicate that cDCs and pDCs are largely generated by progenitors that have already branched.

**Figure 2.**
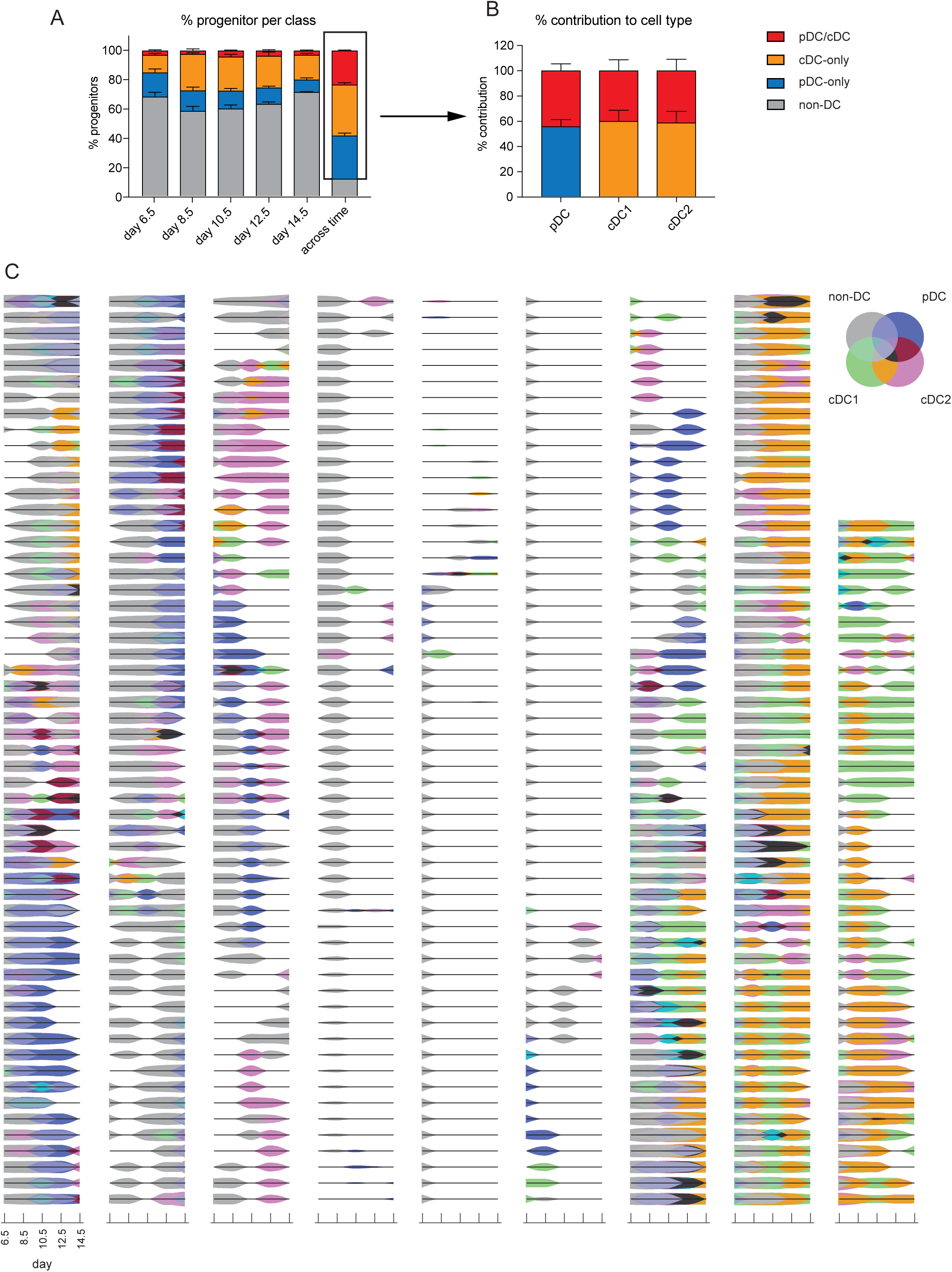
Longitudinal tracking allows accurate interpretation of clonal fate. (A) Classification of clones into 4 classes including non-DC, pDC-only, cDC-only and pDC/cDC based on subtype output at a single time point or across time. (B): % contribution to cell types from three classes of clones based on across time definition. (C) Violin plots showing clonal output of individual barcodes over time. The width of the violin is proportional to the contribution of the clone to the corresponding cell type at that time point. Data shown in A & B are average + SEM of 3 independent cultures from one experiment, representative of 3 independent experiments. C shows the same clones as in Figure 1D & E, from one representative culture.

### Di-SNE movies allow visualization of clonal dynamics

While heatmaps and correlation matrices are useful for a static summary of time-series data for multiple cell types, interpretation of the kinetics of clonal contribution is difficult. Therefore, we utilized the dimensionality reduction technique t-Distributed Stochastic Neighborhood Embedding (t-SNE) (Van der Maaten and Hinton, 2008) to better facilitate visualization of this multidimensional data set. The properties of clones in terms of subtypes and number of cells produced at different times dictated the position of each barcoded HSPC. To visualize clonal fate and DC biomass, we created ‘t-SNE pie maps’ by generating a pie chart representing the proportional output to different DC subtypes and altering point size, respectively (Figure 3). Finally, we linked these series of t-SNE pie maps in the form of a movie for dynamic visualization (Movie S1-3) where changes in point size were interpolated between flanking data points during DC development. We therefore term these developmental interpolated t-SNE (Di-SNE) movies.

**Figure 3.**
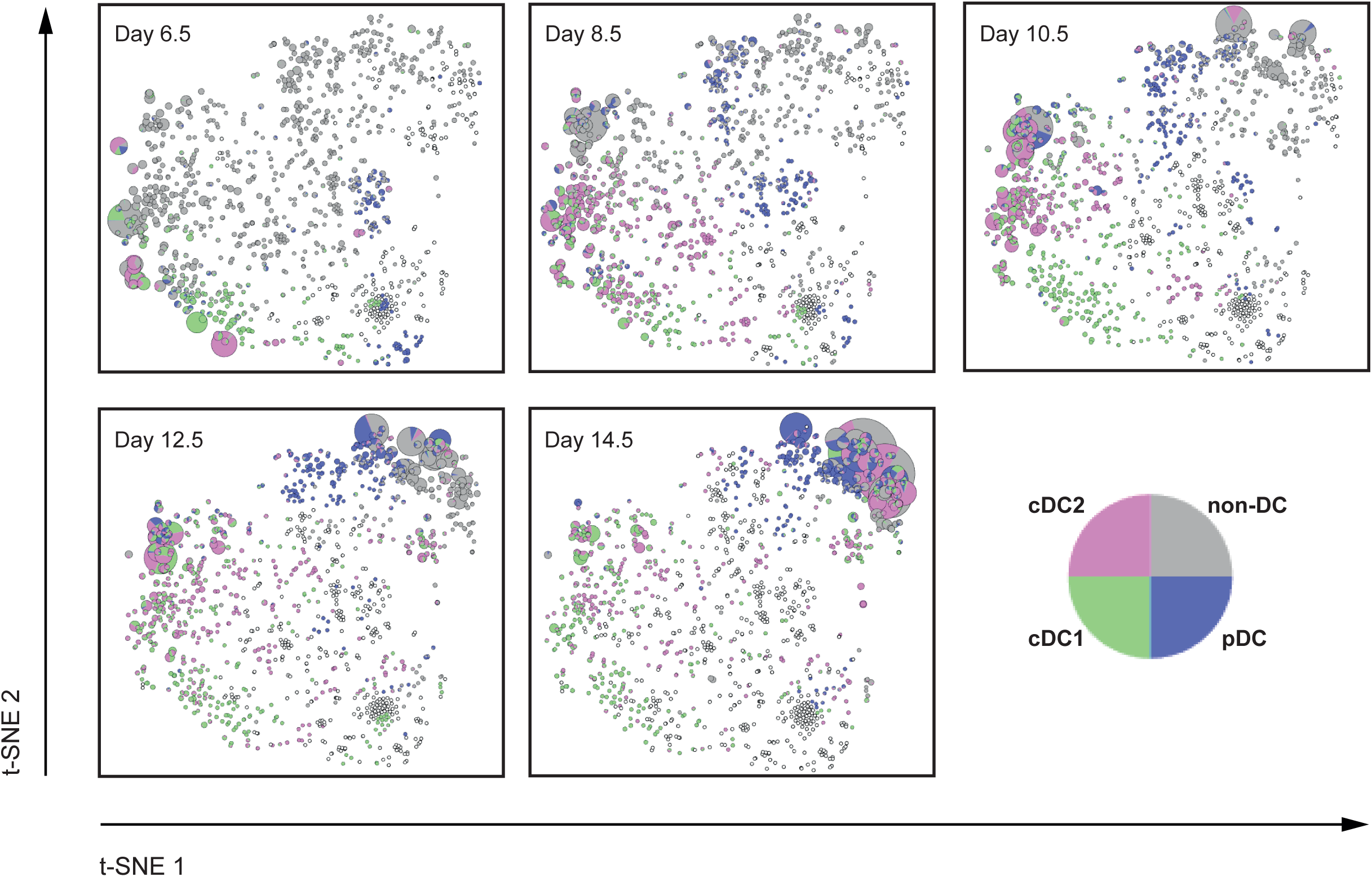
Visualizing diversity of DC cellular trajectories using Di-SNE. Static t-SNE piemap at each time point (see Supplementary Movie 1 for a dynamic visualization). Each circle represents a barcode-labeled progenitor and the size scaled to the number of cells produced by that clone per time point. Each sector in the pie chart represents the proportion of each cell type produced. Data are pooled from three independent cultures (368 data points each (out of 368, 410 and 384) for equivalence) from one experiment, representative of 3 independent experiments. Supplementary Movies 2 & 3 show results from the other 2 experiments.

We performed Di-SNE visualization on data pooled from three independent wells, incorporating all time points available (Figure 3, Movie S1). Similar to the heatmap representation, heterogeneity was observed but patterns were more easily distinguishable considering the bias was incorporated into one pie, rather than four elements, and that clone size was better represented through dot size rather than color. These Di-SNE movies (Movie S1-3) portrayed the dynamic process of DC development encompassing the complexities of qualitative, quantitative, and now temporal characteristics of each clone underlying development. Therefore, Di-SNE movies are an effective and powerful tool for visualization of clonal dynamics, and this technique has been packaged into a stand-alone software package *PieMaker* (supplementary file 3).

### Multiple trajectories of DC development

To further characterize the clonal dynamics of DC development, we compared several clustering methods and found them to give similar results (Figure S3). We then used the clustering method that was most coherent with our Di-SNE visualization. Specifically, we applied Density-based spatial clustering of applications with noise (DBSCAN) (Ester et al., 1996) on the t-SNE map to identify 16 clusters (Figure 4A). Visualization using spindle plots for each cluster showed that clusters were mainly separated by distinct fate bias or timing of contribution with similar fate output (Figure 4B). Interestingly, t-SNE mostly positioned clusters with a similar fate but asynchronous waves of contribution across a band in the plot (see manually annotated circles in Figure 4A) to form four major groups of trajectories including cDC-biased, pDC-biased, multipotent and a group of very small clones with mixed output (Figure 4B&F). There was large variation in the number of clones and DC biomass produced by each cluster (Figure 4C). The most prominent trajectory was cDC-biased, which comprised of ∼43% of clones that contributed ∼60% of cDC generation (Figure 4D&E). Similarly, ∼33% of clones followed a pDC-biased trajectory, which generated more than half of pDCs (Figure 4D&E). Only 12% of clones were identified in the multipotent clusters, which contributed to 36% of pDCs, 31% of cDC1s and 39% of cDC2s (Figure 4D&E). In addition, cluster 2 was in a region containing very small clones that were mostly unipotent. These represented 12% of total clones and less than 1% of total number of DCs generated (Figure 4D&E). Importantly, independent wells within the same experiment were reproducible by comparing the occurrence of barcodes in each cluster (Figure 4G), and between experiments using Jensen-Shannon (JS) divergence (Amir et al., 2013) to holistically assess similarity between data sets (Figure 4H). Thus, we have identified multiple major trajectories of DC development and demonstrated the majority of clones within the HSPC fraction, but not all, follow cDC- or pDC-biased trajectories that contribute to the majority of their biomass.

**Figure 4.**
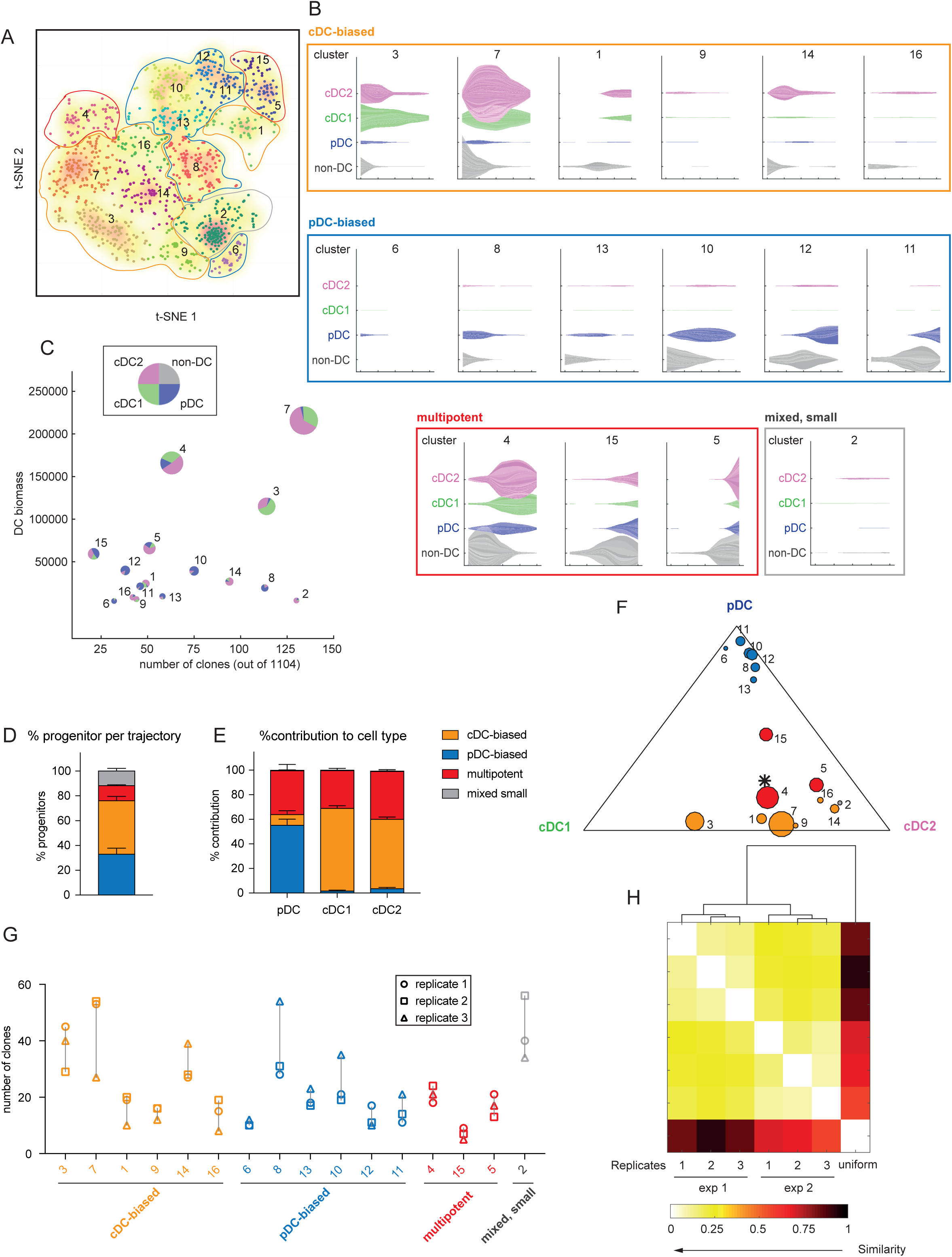
Identification of major classes of DC cellular trajectories. (A) DBSCAN-based algorithm identifies 16 clusters on the t-SNE map (as in Figure 3). Most clusters correlate well with the overlaid barcode density heatmap. The clusters are manually annotated into 4 major classes of trajectories based on distinct fate output. (B) Spindle plots showing contribution to each subtype over time by clones from individual clusters. The width of the spindle is proportional to the contribution of the cluster to the corresponding cell type at that time point, and each partition of the spindle (varying colour shades) represents individual clones within the cluster. (C) Each cluster is quantified both in terms of the number of clones (out of a total of 1104, pool of 3 independent cultures) it includes (x-axis) and DC biomass (the number of DCs it contributes) (y-axis, pie radius). Pie charts show cluster compositions in terms of DC subtypes. (D) % progenitors from each trajectory class as defined in (A & B). (E) % contribution to cell types by each trajectory class. D&E: average + SEM of 3 independent cultures. (F) Ternary plot showing subtype bias of each cluster. Circle size is proportional to DC biomass of the cluster. Asterisk denotes the population average. (G) Barcode representation from the 3 independent cultures in each cluster. (H) Jensen-Shannon (JS) divergence measuring the similarity between independent cultures within the same experiment (very low value, highly similar pattern); between 2 independent experiments (low value, reproducible pattern); and between uniformly distributed pattern on the defined t-SNE region (high value, dissimilar pattern).

### Cellular trajectories are intrinsically programmed

Next, we asked whether the cellular trajectories of members of a single clone are highly correlated i.e. were they programed for this trajectory? To this end, we applied clone-splitting by first pre-expanding barcoded progenitors for 4.5 days and then equally split the wells into two parallel FL cultures (Figure 5A). We then performed serial sampling and barcode analysis on both arms of the experiment as described. We compared the fate and clone size of shared barcodes between replicates (58% in experiment 1 & 73% in experiment 2) in parallel cultures across all time points (Figure 5 & S4). Fate conservation was defined using JS divergence or cosine similarity, where both measured similarity in clonal kinetics (types of progeny produced and the order) and produced similar results (Figure S4B). Size conservation was measured as the base two logarithm of ratio of biomass between the shared barcodes, which essentially measured the discrepancy in division number between splits. Interestingly, we found that many clones were concordant in their cellular trajectories between parallel cultures implying descendant cells carried a ‘memory’ of what DCs to make, when to make them, when to change fate output and how many cells to produce (Figure 5 & S4A). These results are consistent with fate being a heterogeneous, yet intrinsic and heritable property of individual founder cells when measured in similar environments.

**Figure 5.**
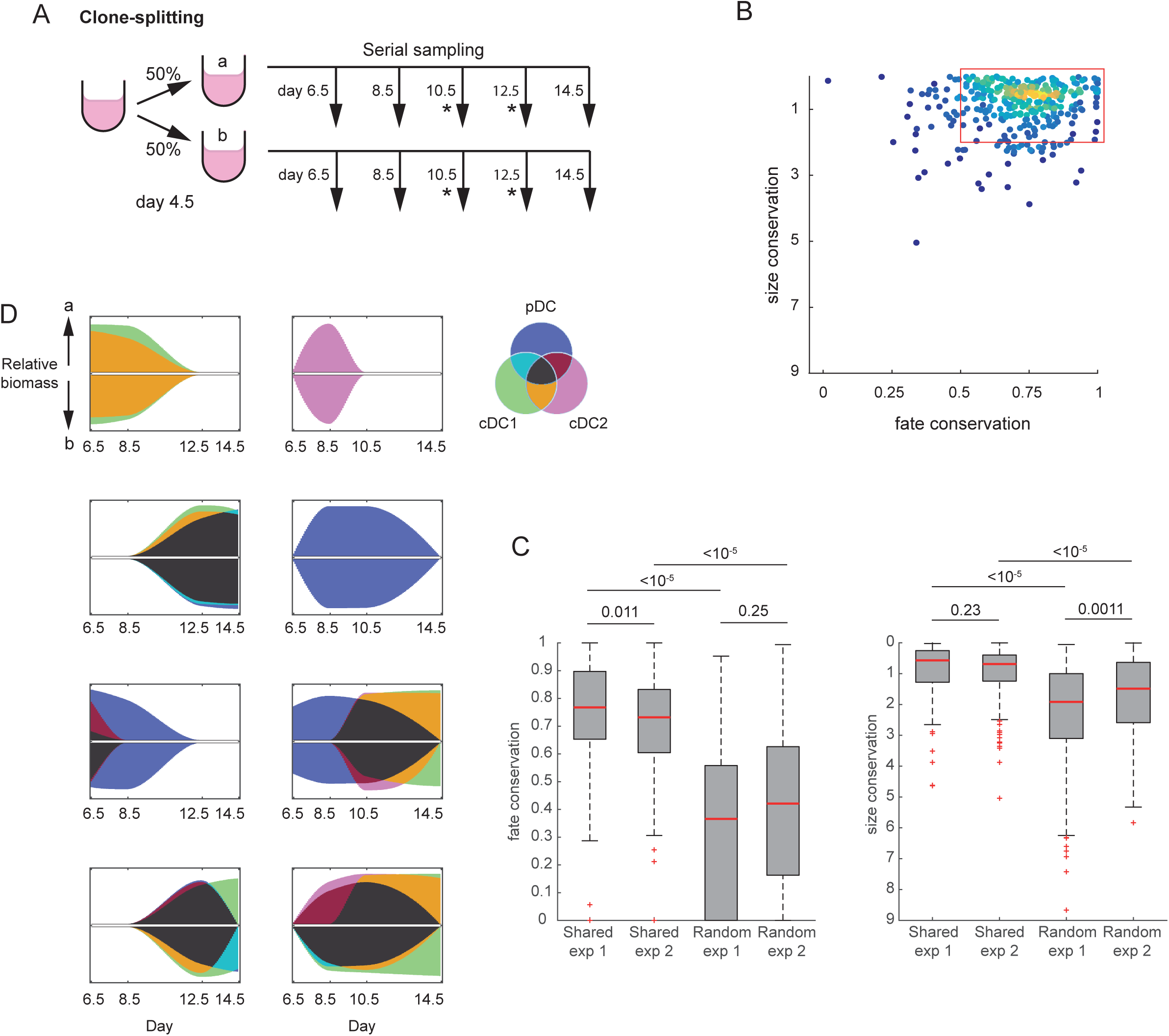
Clonal cellular trajectories are largely programmed. (A) Schematic of clone splitting experiment. Barcoded progenitors were pre-expanded in FL culture for 4.5 days and split into two parallel cultures (a & b). Serial sampling was then performed from both arms as described in Figure 1A. *: data from day 10.5 is lacking in experiment 1 and day 12.5 is lacking in experiment 2 due to technical issues. (B) Conservation of shared barcodes across all time points. Each point represents a barcode with reads detected in both halves of the split culture. For each barcode, size conservation is defined as the base two logarithm of ratio of total read counts, and fate conservation is defined as JS divergence. Clones inside the gate represent 80% of total shared barcodes, which contributes to 80% of total biomass. Data are a pool of 2 sets of parallel cultures from experiment 1, representative of 2 independent experiments. (C) Summary of fate and clone conservation value comparing split barcodes with randomly paired unrelated barcodes. Boxplots span interquartile range. Pooled data from both independent experiments are shown. Statistical significance is measured using Mann-Whitney U test. (D) Paired violin plots comparing cellular trajectories from two arms of split culture (a vs b). Eight examples of clones with high conservation values are shown. Full list of clones from experiment 1 is shown in Figure S4.

## DISCUSSION

The framework developed here provides a statistically robust, quantitative, visually intuitive approach for the tracking of thousands of clones over short periods of time with separation into clusters. It allows systematic examination of lineage trajectories of any developmental system, where cells can be cultured *ex vivo* and subsampled at desired time intervals. Our results indicate that assessment of *bona fide* clonal cellular trajectories is crucial to accurately determine clonal fate, as opposed to measuring fate at a fixed time point. In addition, by incorporating clone-splitting, we demonstrate that clonal fate and waves of contribution to DCs is heterogeneous yet largely programmed early in the developing clone. This provides the rationale to combine our method with other approaches such as single cell RNA-seq in parallel to not only measure cellular trajectories but the underlying molecular trajectories that guide these properties, or to test the effect of biological variation or perturbation such as drug treatment and genetic manipulation on one arm of the clone splitting experiment.

Importantly, we demonstrate that the majority of HSPCs already have a cDC- or pDC-biased fate by measuring clonal output across multiple time points. Our results do not support the current model, which implies a common origin of cDCs and pDCs from CDPs (Guilliams et al., 2014b). This could be partly explained in that many prior studies do not incorporate CCR9 to define pDCs, leading to possible misallocation of cDC precursors as pDCs. Similarly, two recent studies (See et al., 2017; Villani et al., 2017) using single cell profiling of the human DC compartment independently identify contaminating DC precursors within the phenotypically defined pDCs. Future studies should determine whether those populations and murine SiglecH^+^CCR9^−^ cells represent the same precursor population. Our observation of early cDC and pDC bifurcation is also partly supported by the identification of cDC-, cDC1- and cDC2-committed progenitors in various fractions of HSPCs and downstream progenitors (Grajales-Reyes et al., 2015; Schlitzer et al., 2015; Schraml et al., 2013). Importantly, our results indicate that a pDC-committed progenitor population likely exists within HSPC fraction, indicative of early branching, similar to a recent study (Lee et al., 2017).

Our results highlight a remarkable degree of heterogeneity within early HSPC population. Longer term efforts should appraise not only progenitors but also their progeny at a single cell level to determine how origin dictates functional heterogeneity. This information, combined with the molecular drivers that underlie true cellular trajectories, and within an *in vivo* context, are necessary for a full understanding of development.

## AUTHOR CONTRIBUTIONS

Conceptualization, D.L, A.K. J.G. and S.H.N; Methodology, D.L., A.K., J.G. and S.H.N.; Software, A.K. and J.G.; Formal Analysis, A.K.; Investigation, D.L.; Visualization, D.L. and A.K.; Writing Original Draft, D.L., A.K. and S.H.N.; Writing – Review & Editing, D.L., A.K., J.G, E.C., P.D.H. and S.H.N.; Supervision, S.H.N., E.C. and P.D.H.

### ACKNOWLEDGMENTS

We thank the WEHI FACS laboratory, Dr. Stephen Wilcox and Dr. Tom Weber for technical support. We thank Dr. Samir Taoudi, Prof. Gabrielle Belz and Prof. Stephen Nutt for insightful discussions. This work was supported by the National Health & Medical Research Council, Australian Government Grant APP1100033.

## METHODS

### Mice

All mice were bred and maintained under specific pathogen-free conditions at WEHI, according to institutional guidelines. Either C57BL/6 (CD45.2) or C57BL/6 Pep^3b^ (CD45.1) male mice aged between 8-16 weeks were used.

### Isolation of bone marrow progenitors

Bone marrow cells from hip, tibia and femur were stained with anti-CD117 allophycocyanin (APC) at 4 ^°^C for at least 30 minutes. CD117 enrichment was then performed using MACS after incubation with anti-APC magnetic beads according to manufacturer’s protocol (Miltenyl Biotec). CD117-enriched cells were stained with anti-Sca1 antibody. Finally, cells were resuspended in FACS buffer (PBS with 0.5% FBS and 2 mM EDTA) containing propidium iodide (1 μg/ml) before sorting cKit^+^Sca1^+^ (SK) cells using a BD Influx, BD Fusion or BD FACSAria-II/III (Beckton Dickinson).

### Barcode transduction

Barcode transduction was performed as described (Naik et al., 2013). Briefly, progenitors were transferred into a 96-well round bottom plate at less than 1 x 10^5^ cells/well in 100 μl StemSpan medium (Stem Cell Technologies) with 50 ng/ml stem cell factor (generated in-house) and small amount of lentivirus containing the barcode library and GFP reporter (Naik et al., 2013). The amount of lentivirus was pre-determined in control experiments to give approximately 10% transduction efficiency. The plate was centrifuged at 900 *g* for 90 minutes at 22 °C prior to incubation at 37 °C and 5% CO_2_-in-air for 4.5 hours. After transduction, cells were washed using a large volume of RPMI containing 10% FBS and resuspended in FL-supplemented DC conditioned medium (Naik et al., 2005).

### FL culture and serial sampling

5×10^3^ labeled cells were cultured with 200 μl DC conditioned medium supplemented with hFL (BioXcell, 800 ng/ml) per well in a 96-well round-bottom plate. After 6.5 days of culture, cells were gently mixed a few times with a pipette and half were removed for subtype isolation and another half were kept in culture with medium topped up to 200 μl to allow further DC development. Three mature DC subtypes and non-DCs (other live cells) were sorted. The same procedure was repeated every two days. At the last time point, all cells from each well were harvested and sorted.

Sampling controls were included to test whether there was an equal chance of capturing the same barcodes in two fractions. To control for the first time point, wells were split and each half was analyzed by sorting DC subtypes and recovering barcodes. To control for the second time point, half of culture was discarded at day 6.5 and the other half was kept in wells and then each half was analyzed at day 8.5. Similarly, to control for later time points, half of culture were discarded and the other half were kept in wells at each time point prior to analysis.

Clone-splitting experiments were performed to assess conservation of fate between shared barcodes over time. Briefly, wells were split into two at day 4.5 and both cultured until day 6.5. After that, serial sampling every two days and analysis was performed from each split well as the other samples.

### Isolation of DC subtypes

Cells removed from culture were stained with antibodies against CD11c, MHCII, Siglec-H, CCR9, Sirpα and CD24. pDCs were gated as CD11c^+^MHCII^-/low^Siglec-H^+^CCR9^+^. cDC1s were gated as CD11c^+^MHCII^+^Sirpα^−^CD24^hi^. cDC2s were gated as CD11c^+^MHCII^+^Sirpα^+^CD24^+^. Fractions other than these three DC subtypes were sorted together as non-DCs.

### Barcode amplification

PCR and sequencing were performed as described previously (Naik et al., 2013). Briefly, sorted populations were lysed in 40 μl Viagen lysis buffer containing 0.5 mg/ml Proteinase K (Invitrogen) and split into technical replicates. Barcodes in cell lysate were then amplified following two rounds of PCRs. The first PCR amplified barcode DNA using a common primer, and second PCR introduced a sequencing primer with flow cell attachment and a well-specific index primer to each sample for later de-multiplexing *in silico*. Caution was taken at all steps to avoid barcode contamination between samples. Products from second round PCR with index primers were run on a 2% agarose gel to confirm a PCR product was generated, prior to being cleaned with size selected beads (NucleoMag NGS) according to the manufacturer’s protocol. The cleaned PCR products were pooled and deep sequencing was performed on the Illumina NextSeq platform.

### Processing of sequencing data

Barcode data processing was performed largely as described previously (Naik et al., 2013). First, number of reads per barcode in each technical replicate from each sorted sample was calculated using processAmplicons function from edgeR package (Robinson et al., 2010). The quality of the samples was assessed by comparing technical replicates. The majority of samples used across all experiments (253 out of 264) met the following criteria: (1) average total read number across the two replicates was ≥ 10^4^; (2) ratio between the smaller and the larger number of total reads was ≥ 0.2 (i.e. less than one order of magnitude); and (3) Pearson correlation coefficient between barcode read counts in two replicates was ≥ 0.6.

Read counts between technical replicates from the samples were averaged, except for barcodes that had reads in one technical replicate but not the other due to technical reasons, which were set to zero read count. Total read counts of each sample was then normalized to 10^5^.

### Barcode categorization

Each barcode was categorized into 4 classes: “non-DC” (produced no DCs), “pDC-only” (produced pDCs, but neither of cDC1 or cDC2), “cDC-only” (produced cDC1 and/or cDC2 cells, but no pDC) and “pDC/cDC” (produced pDCs and cells of either one or both cDC1 and cDC2 types). To assign a particular DC subtype fate to a barcode, two parameters were used including minimal read count and minimal proportion. In Figure 2A, minimal read count was set to 750 and minimal proportion was set to 5%. For example, if barcode A had 1,000 reads in pDC, 600 reads in cDC1 and 90,000 reads in cDC2, the cDC1 reads was first set to zero as it did not pass the minimal read count threshold. As this barcode produced 1% pDC (< 5%) and 99% cDC2, it was classified as cDC-only clone. Categorization was performed based on data at each time point independently, or based on data across time. For categorization across time, a barcode was considered to produce a certain subtype X if it produced subtype X at any of the time points. Categorization was repeated using varying value combination of the two thresholds to verify that small changes in the values of these parameters qualitatively results in same outcomes (Figure S1E).

### Biomass computation

Since every sample (e.g., “pDC at day 6.5” or “cDC1 at day 10.5”) is normalized to 10^#^ reads, barcode read counts only reflect the contribution of the barcode to a particular subtype but not to the entire culture. Therefore, barcode biomass per time point was computed using normalized barcode read counts (as described above) and subtype proportion (number of cells per subtype / sum of all subtypes) as inputs, so that the sum of all barcode biomass at any time point represents 100% of the culture.

### Heatmaps

Heatmaps were generated to visualize clonal output to DC subtypes at different time points using barcode biomass multiplied by a factor of 100 and hyperbolic arcsine transformed. Such a transformation resembles log-transformation with pre-selected logarithm base. The advantage of using hyperbolic arcsine is that this function is defined at zero. The order of barcodes in both visualization methods was produced using an algorithm for optimal leaf ordering for hierarchical clustering (Bar-Joseph et al., 2001), as implemented in optimalleaforder function from MATLAB R2015b (MathWorks). In brief, such an ordering maximizes sum of similarities between adjacent barcodes, e.g., adjacent rows in the heatmap.

### Visualization using t-SNE, static pie maps and Di-SNE movie

First, t-distributed stochastic neighboring embedding method (t-SNE) was performed with default parameters to reduce the dimensions of the dataset to 2D (Van der Maaten and Hinton, 2008). Hyperbolic arcsine-transformed biomass counts from all time points were pooled from three independent cultures as input to t-SNE algorithm. Barcodes that did not produce any DCs were excluded. The output of t-SNE was used for downstream visualization and clustering. To visualize clonal output and size on the t-SNE map, each barcode was represented as a pie chart (t-SNE pie maps). The segments of the chart depict the proportion of each subtype at a particular time point. The radius of the pie chart reflects the total biomass of the given barcode at the given time point. For the purpose of visualizing individual cellular trajectories (developmental changes over time) and clusters (see below), a cubic spline based interpolation for time values between experimental time points was applied. Depending on the settings for Di-SNE movie generation, linearly interpolated frames can be added between frames that correspond to experimental time points (see manual for PieMaker software).

### Clustering

To identify major patterns, several clustering methods were applied including Density-Based Spatial Clustering of Applications with Noise (DBSCAN) algorithm (Ester et al., 1996), Gaussian-mixture clustering and Affinity Propagation algorithm (Frey and Dueck, 2007). These methods were applied on both raw data and using scatter plots derived from t-SNE as input. These methods were capable of producing similar results. For example, raw data based clustering resulted in clusters that were spatially consistent when projected onto t-SNE plots (Figure S3A) and produced trajectories with similar patterns (data not shown).

A barcode density plot using kernel density estimation via diffusion (Botev et al., 2010) was generated to assess feasibility of each particular clustering method by first running each of the three algorithms on grids of parameter values, and visually inspecting how well the resulting clustering aligned with the barcode density plot. DBSCAN-based clustering was found to align with the density plot best. Therefore, DBSCAN was used to identify cluster centroids, and each unassigned point was assigned to the cluster with nearest centroids. The resulting clusters were manually categorized into “cDC-biased”, “pDC-biased”, “multipotent” or “mixed, small” based on visual inspection of the corresponding Di-SNE movie, visualization of subtype output per cluster using spindle plots (Figure 3B) and visualization of fate bias per cluster using ternary plots (Figure 3F). Spindle plots showed the total biomass per DC subtypes for all barcodes that are members of that cluster. The spindle plots are stacks of biomasses of barcodes included in the corresponding cluster. Individual barcodes can be distinguished by varying color shades. The ternary plot was generated using proportions of pDC, cDC1 and cDC2 biomasses (non-DC excluded) to define coordinates for each cluster in the equilateral triangles.

### Conservation between shared barcodes in split cultures

Given a split culture, barcodes without any DC biomass in each of the split parts (non-DCs only) and barcodes that were present only in one of the splits were excluded and the rest were identified as shared barcodes. Fate conservation was computed to measure similarity between kinetics of DC subtype production (e.g., whether split parts of the same clone produced same types of DC and in the same order). First, biomass values were hyperbolic arcsine-transformed and Jensen-Shannon (JS) divergence and cosine similarity were computed. Both methods produced very similar results, and hence JS divergence was used to estimate fate conservation in Figure 5. Size conservation was computed to measure similarity in clonal expansion between the split parts of the same clone. First, total biomass per barcode was calculated (sum of biomass from all subtypes from all time points) for each split part. Next, ratio of the smaller total biomass to the larger was calculated and base two logarithm of this ratio was computed as a measure of size conservation. For example, a difference of one could be interpreted that one part of the clone made on average one more division round. Note that biomass of non-DCs were excluded during computation of both fate and size conservation. Random controls were generated by randomly paired unrelated barcodes in the same culture to assess whether the observed conservation was due to chance.

### Statistical analysis

Mann-Whitney U test was performed to measure the significance of the observed difference between groups. All data were presented in boxplots that span interquartile range.

### Data and codes availability statement

All data and codes that support the findings of the study are included in the supplementary files.

---

**Figure S1.**
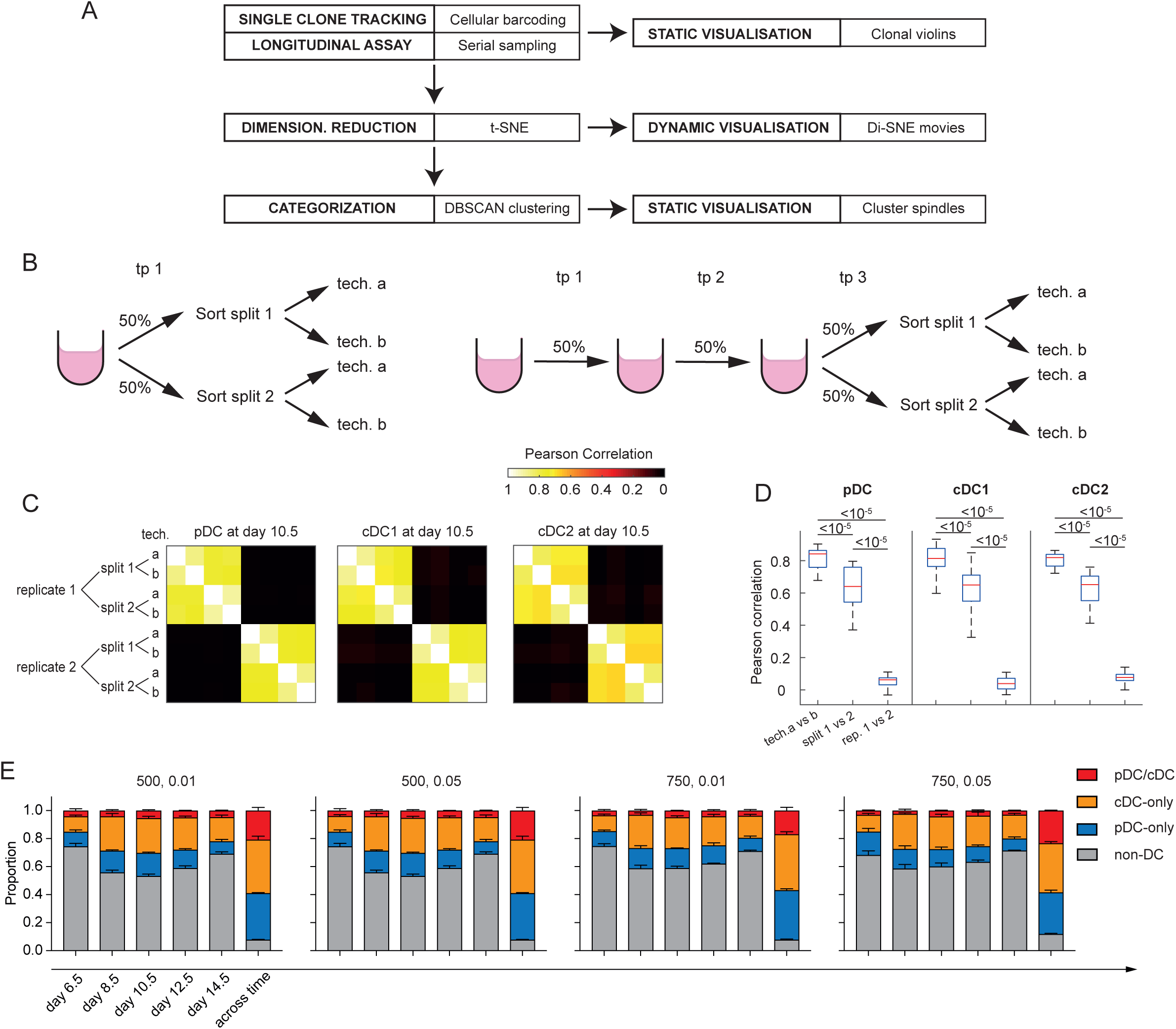
Optimization and validation of experimental set up. (A) Workflow used in this framework to study clonal dynamics. Left boxes: general methods; Right boxes: specific methods used in the study. (B) Schematic of sampling controls. Same sampling procedure was performed as described in Figure 1A until the corresponding time point when each split was sorted separately and barcode signatures were compared. (C) Correlation heatmaps comparing Pearson Correlation between technical replicates (tech a vs b, split after sorting), sampling controls (split 1 vs 2) and unrelated samples (replicate 1 vs 2). Examples from day 10.5 are shown. Pearson correlation was calculated using hyperbolic arcsine-transformed values of read counts before pre-precessing. (D) Summary of Pearson correlation of all samples at all time points from one experiment. Boxplots show interquartile range; middle bar depicts median. One experiment was performed, with two replicates per time point. Mann-Whitney U test was performed. (E) Barcode categorization using different thresholds (minimal read counts, minimal proportion) as described in methods.

**Figure S2.**
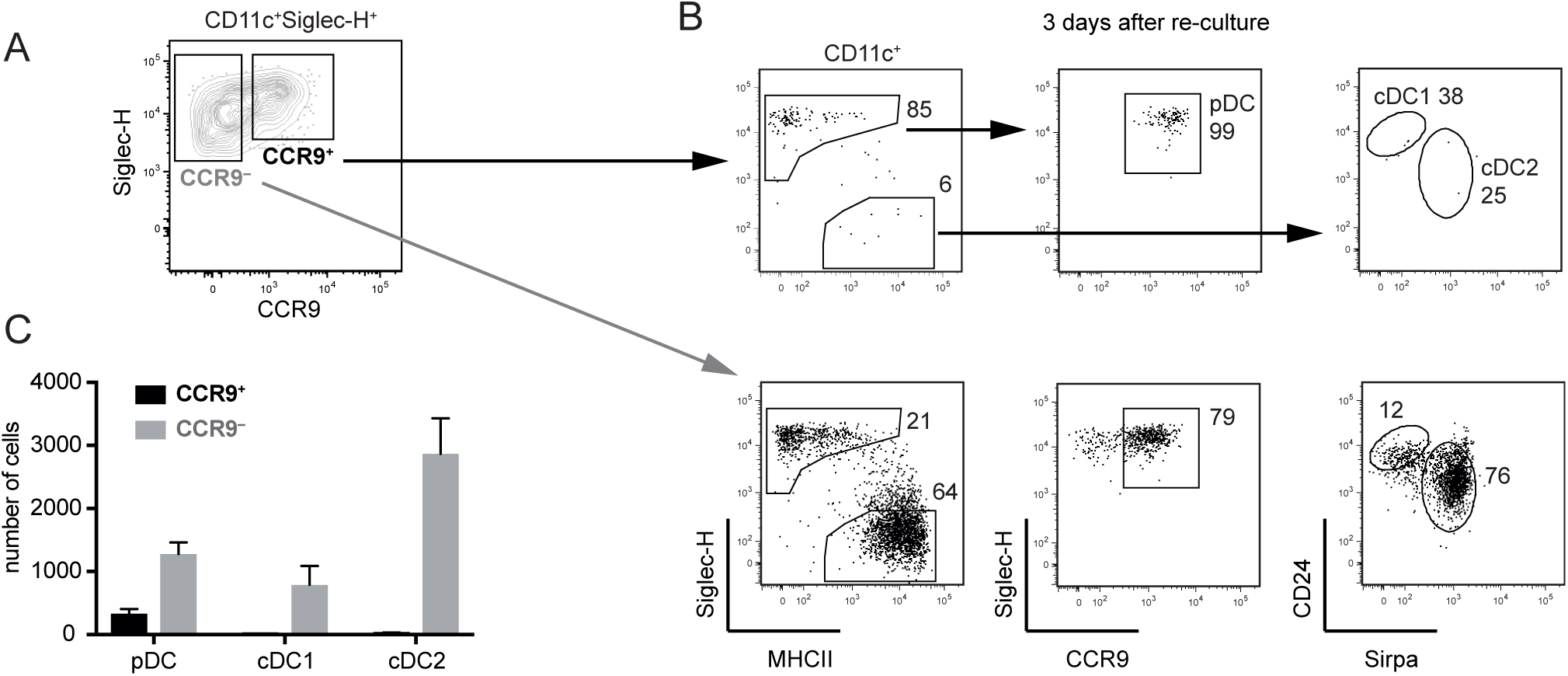
Siglec-H^+^CCR9^−^ cells are DC precursors. (A) CCR9^+^ and CCR9^−^ cells from the CD11c^+^Siglec-H^+^ fraction of cultured DCs were sorted. (B) Cells were re-cultured and analyzed by FACS at day 3. CCR9^−^ cells generated both cDCs (Siglec-H^−^MHCII^hi^) and pDCs (Siglec-H^+^CCR9^+^) upon re-culture, while viable CCR9^+^ cells maintained the original phenotype (Siglec-H^+^CCR9^+^). Numbers in example FACS plot depict percentage of parent gate. (C) Summary of number of cells generated at day 3 from Siglec-H^+^CCR9^+^ and Siglec-H^+^CCR9^−^ cells. Data shown are average + SEM of 3 replicates, representative of 3 independent experiments.

**Figure S3.**
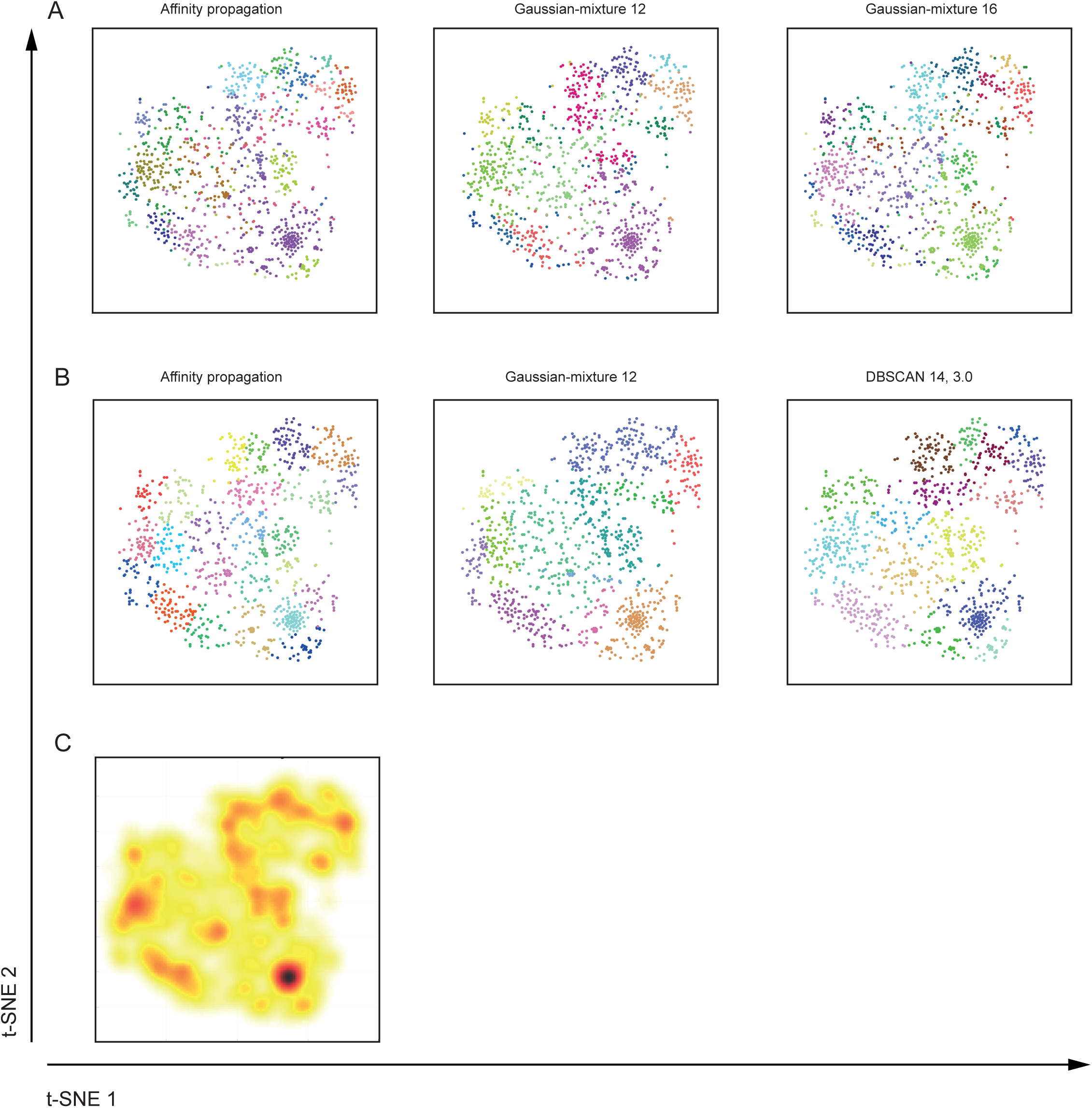
Comparing different clustering methods. (A) Clustering using raw data as input. Clusters produced by affinity propagation and Gaussian-mixture clustering (12 or 16 components) are projected onto t-SNE plots. Most clusters produced are spatially consistent with t-SNE. (B) Clustering using t-SNE as input. Clusters produced by affinity propagation, Gaussian-mixture clustering (12 components) and DBSCAN (same as Figure 4A) are shown. Note that DBSCAN produces clusters that correlate best with the dense regions of barcodes in (C) t-SNE density heatmap.

**Figure S4.**
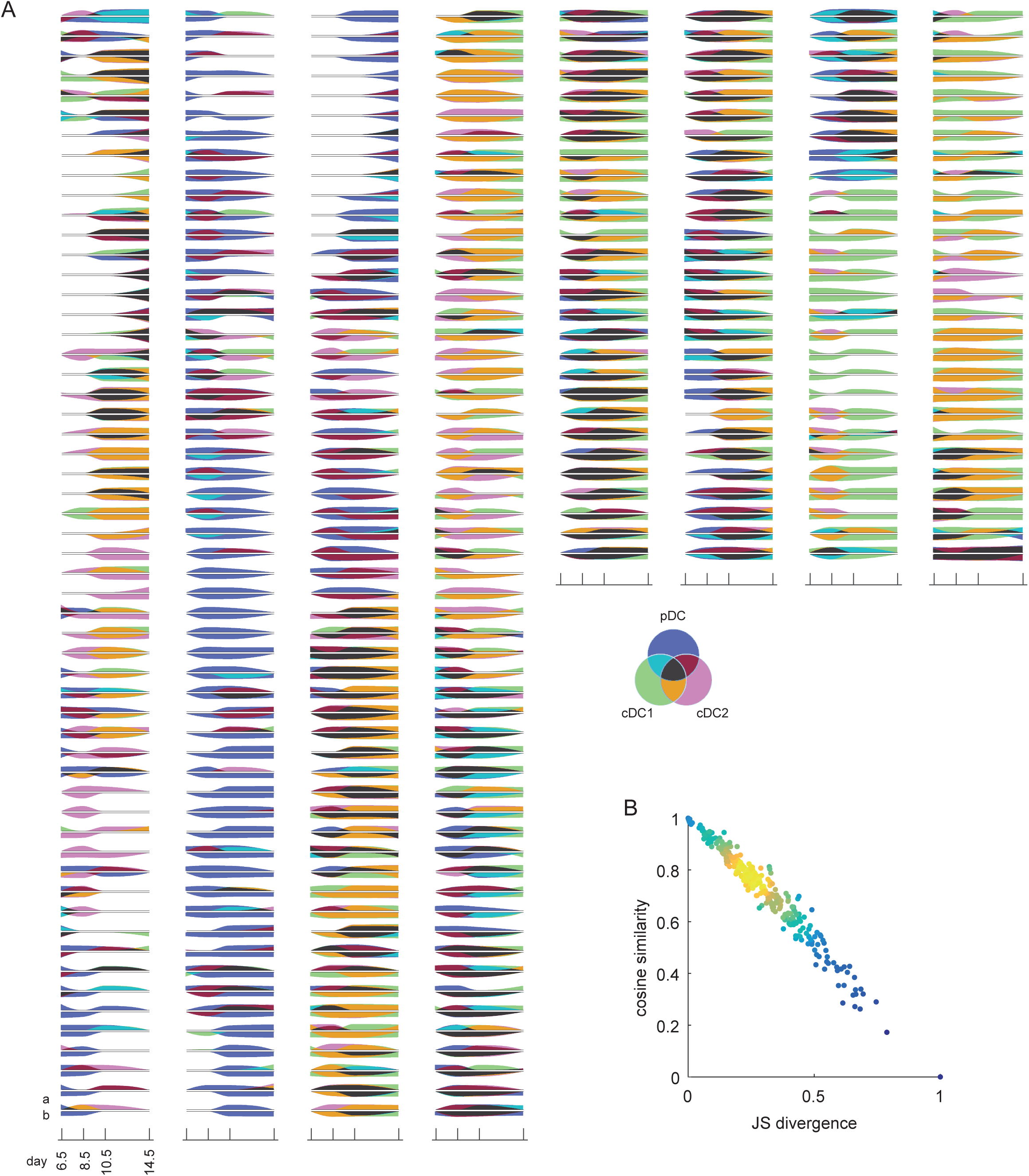
Comparison of cellular trajectories in clone split cultures. (A) Full list of paired violin plots comparing trajectories of shared barcodes (a vs b) as in Figure 5D. (B) Linear correlation between JS divergence and cosine similarity to measure fate conservation of shared barcodes. Cosine similarity of 1 or JS divergence of 0 indicates identical output. Each dot represents one shared barcode. Color depicts density of barcodes. Data are pooled from 2 sets of parallel cultures from experiment 1, representative of 2 independent experiments.

